# Calcium transients trigger switch-like discharge of prostaglandin E_2_ in an extracellular signal-regulated kinase-dependent manner

**DOI:** 10.1101/2023.02.01.526734

**Authors:** Tetsuya Watabe, Shinya Yamahira, Kanako Takakura, Dean Thumkeo, Shuh Narumiya, Michiyuki Matsuda, Kenta Terai

## Abstract

Prostaglandin E_2_ (PGE_2_) is a key player in a plethora of physiological and pathological events. Nevertheless, little is known about the dynamics of PGE_2_ secretion from a single cell and its effect on the neighboring cells. Here, by observing confluent Madin-Darby canine kidney (MDCK) epithelial cells expressing fluorescent biosensors we demonstrate that calcium transients in a single cell cause PGE_2_-mediated radial spread of PKA activation (RSPA) in neighboring cells. By *in vivo* imaging, RSPA was also observed in the basal layer of the mouse epidermis. Experiments with an optogenetic tool revealed a switch-like PGE_2_ discharge in response to the increasing cytoplasmic Ca^2+^ concentrations. The cell density of MDCK cells correlated with the frequencies of calcium transients and the following RSPA. The extracellular signal-regulated kinase (ERK) activation also enhanced the frequency of RSPA in MDCK and *in vivo*. Thus, the PGE_2_ discharge is regulated temporally by calcium transients and ERK activity.

## Introduction

Prostaglandin E_2_ (PGE_2_) is an eicosanoid lipid mediator that regulates a plethora of homeostatic functions including vascular permeability, immune response, and mucosal integrity (Narumiya, 2007). The metabolic pathway of PGE_2_ production, which is a major branch of the arachidonic acid cascade, was extensively studied in the mid to late 20th century. This cascade starts from the activation of cytosolic phospholipase A2 (cPLA2) (Park et al., 2006). Increased intracellular calcium induces the translocation and activation of cPLA2 from the cytosol to the Golgi, endoplasmic reticulum, and perinuclear membrane, where cPLA2 cleaves arachidonic acid out of the membrane phospholipids (Clark et al., 1995; Evans et al., 2001; Hirabayashi et al., 1999). The arachidonic acid is then presented to cyclooxygenases, COX1 and COX2, to yield PGH_2_, which is further converted to PGE_2_ by prostaglandin E synthases (Smith & Langenbach, 2001). PGE_2_ synthesized *de novo* is secreted to the extracellular space either by passive diffusion or by active transport by multidrug resistance protein 4 (MRP4) (Reid et al., 2003). The secreted PGE_2_ exerts its actions by acting on four G protein-coupled receptors (GPCRs), EP1 to EP4, expressed in neighboring cells (Narumiya, 2007; Regan, 2003).

Although it is largely believed that the PGE_2_ action is primarily regulated by the expression and activation of COX (Kalinski, 2012), cPLA2 appears to play a more important role in the short-term regulation of PGE_2_ production and secretion (Leslie, 2015). It was shown that the secretion of arachidonic acid is induced within a few minutes after the calcium-dependent translocation of cPLA2 to endo-membranes (Evans et al., 2001; Hirabayashi et al., 1999). Moreover, ERK and p38 MAP kinases also contribute to the activation of cPLA2 (Lin et al., 1993), which may be calcium-independent (Gijón et al., 2000). Importantly, these earlier biochemical studies did not elucidate the dynamics of the production and secretion of PGE_2_ at the single-cell level, leaving many questions unanswered. For example, do all cells contribute to the production of PGE_2_? Does each cell keep secreting PGE_2_ upon stimulation?

Genetically encoded biosensors based on the principle of Förster resonance energy transfer (FRET) allow us to visualize the dynamics of intracellular signaling molecules at the single-cell resolution (Greenwald et al., 2019; Miyawaki & Niino, 2015). The development of transgenic mice expressing FRET biosensors has opened a window to the visualization of signaling molecule activity in live tissues (Terai et al., 2019). Furthermore, by using the activation of protein kinase A (PKA) and ERK MAP kinase as surrogate markers, we can also visualize the intercellular communications mediated by Gs-coupled receptors and tyrosine kinase receptors in live tissues (Hino et al., 2020; Konishi et al., 2021). In this study, we show that PGE_2_ discharged from a single cell causes radial spread of PKA activation (RSPA) in neighboring cells in tissue culture and the mouse epidermis. By combining a chemical biology approach, optogenetic stimulation, and a simulation model, we quantitatively analyzed the PGE_2_ secretion and found that the PGE_2_ discharge is regulated temporally by calcium transients and quantitatively by growth factor signaling and cell density.

## Results

### PGE_2_ mediates radial spread of PKA activation (RSPA)

In an attempt to understand intercellular communication under physiologic conditions, we observed the PKA activity of Madin-Darby canine kidney (MDCK) epithelial cells by using the FRET biosensor Booster-PKA (Watabe et al., 2020; Zhang et al., 2001)(Fig. 1A). We noticed that PKA activation propagates from a single cell to neighboring cells under a confluent condition. We named this phenomenon radial spread of PKA activation (RSPA) and pursued underlying mechanisms (Fig. 1B, Video 1). In a typical example, approximately 100 neighboring cells located within a 100 µm distance exhibit firework-like spread of PKA activation, which decays within several minutes. To characterize this phenomenon without preoccupations, we developed a program to identify and characterize RSPA under various conditions (Fig. 1C). The frequency, but not the radius, of RSPA depended on the cell density; i.e., RSPA was observed only when cells were maintained at more than 6×10^4^ cells/cm^2^ (Fig. 1D, 1E). The probability of RSPA in each cell was also increased in a cell density-dependent manner (Fig. S1A). To examine whether the PKA activation correlates with increased intracellular cAMP concentration, we employed another FRET biosensor for cAMP, hyBRET-Epac, and performed a similar experiment (Ponsioen et al., 2004; Watabe et al., 2020)(Fig. S1B, S1C). Although the increment of the FRET ratio was not so remarkable as that of Booster-PKA (Fig. S1D), we found that the pattern of cAMP concentration change is very similar to the activity change of PKA, indicating that a Gs protein-coupled receptor (GsPCR) mediates RSPA (Fig. S1E). This discrepancy between hyBRET-Epac and Booster-PKA may be partially explained by the difference in the dynamic ranges for cAMP signaling in each FRET biosensor (Watabe et al., 2020). Previous transcriptome analysis of MDCK cells showed that ATP receptor and PGE_2_ receptor EP2 are the most abundant GsPCRs (Shukla et al., 2015). Thus, we examined the contribution of the ATP receptor and PGE_2_ receptors EP2 and EP4 by using specific inhibitors (Fig. 1F). Inhibitors against EP2, EP4, and COX, but not the ATPase apyrase, abolished RSPA, indicating that PGE_2_ mediates RSPA.

**Figure 1:**
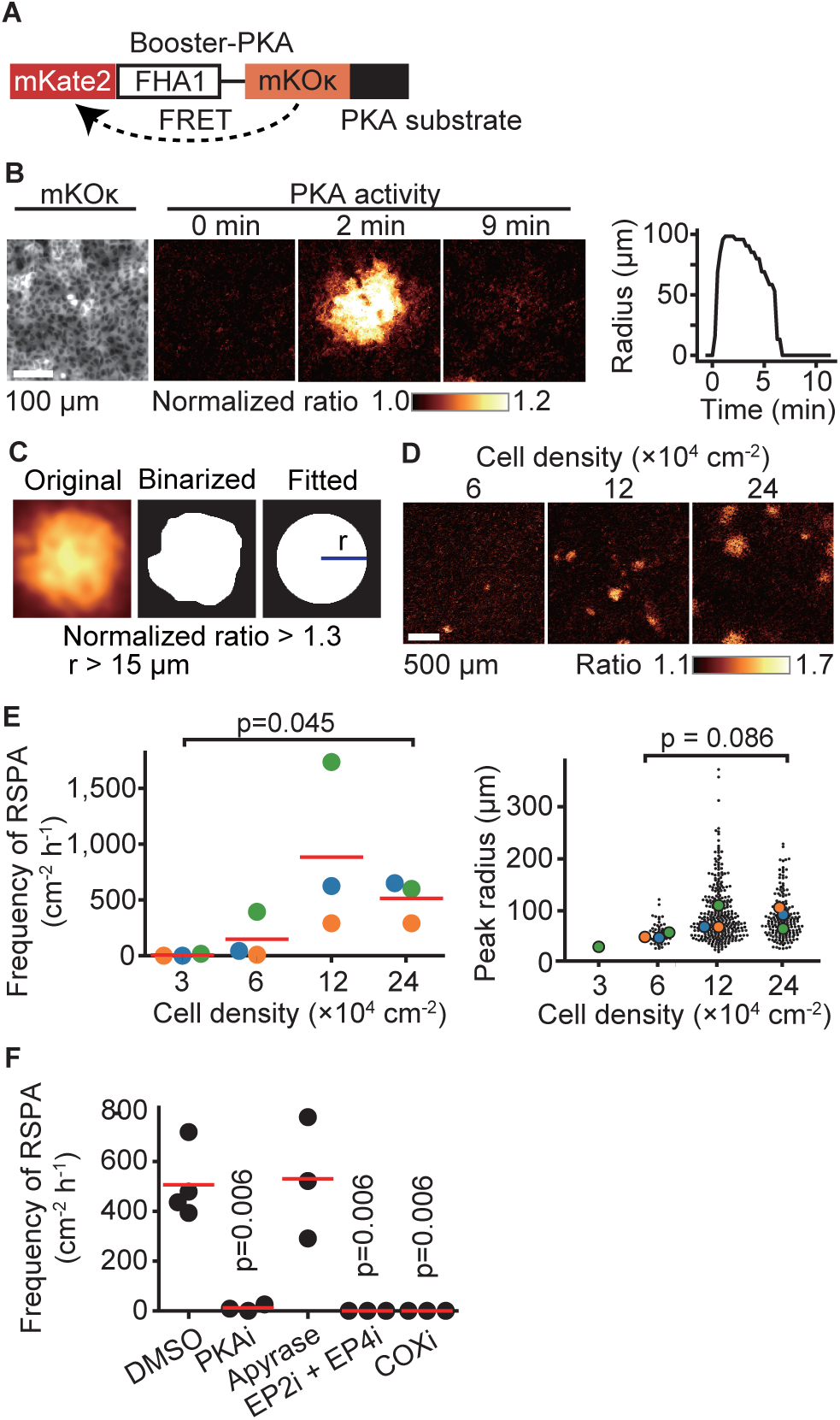
Radial spread of PKA activation, RSPA, in MDCK cells. **(A)** A scheme of Booster-PKA, a PKA sensor. **(B)** Booster-PKA-expressing MDCK cells in a confluent condition were observed every 15 seconds under a fluorescence microscope (Video 1). The image of mKOκ represents the cell density, which is seeded at 2.4 × 10^5^ cells cm-2.mKate2 and mKOκ images were acquired to generate mKate2/mKOκ ratio images representing PKA activity in pseudocolor. The time 0 is set the just before initiation of PKA activation. The radius of RSPA as determined in (C) was plotted as a function of time. **(C)** Procedure to call RSPA positive. The original ratio images were binarized with the threshold value 1.3 of mKate2/mKOκ ratio. The fitted radius of RSPA, r, was defined as the radius of a circle with the same area. When r is greater than 15 µm, it is counted as RSPA. The detailed procedure is in the method section. **(D, E)** MDCK cells expressing Booster-PKA were seeded at the indicated density and analyzed. Representative images in indicated cell densities are shown in pseudocolor (D). Each color in panel E represents an individual experiment. Red lines indicate average values. **(F)** MDCK cells expressing Booster-PKA in the presence of the inhibitors were imaged and analyzed for the RSPA frequency. Reagents are as follows: DMSO, 0.1% v/v DMSO; PKAi, 20 μM H89; Apyrase, 10 unit mL^−1^; EP2i, 10 μM PF-04418948; EP4i, 1 μM ONO-AE3-208; COXi, 10 μM indomethacin. The frequency of RSPA was analyzed 20–80 min after the treatment. Each dot represents an individual experiment. Red lines indicate their average value. p values were calculated between the labeled sample and the DMSO-treated sample.

### RSPA is also observed in the epidermis of the PKAchu mice

To clarify the physiological relevance of RSPA, we used PKAchu mice, which are transgenic mice expressing a FRET biosensor for PKA, AKAR3EV (Kamioka et al., 2012; Sato et al., 2020)(Fig. 2A). We previously observed that ERK MAP kinase activation is propagated radially among the basal layer cells of the mouse epidermis (Hiratsuka et al., 2015), but we failed to observe a similar propagation of PKA activation. We reasoned that this failure was due to the low frequency and short duration of this phenomenon. When we observed a region of over 1 mm square at 1 min intervals, we successfully observed RSPA in the basal layer of the mouse auricular epidermis (Fig. 2B, Video 2). Upon i.p. injection of a COX inhibitor, RSPA almost completely disappeared within 10 min (Fig. 2C), indicating that RSPA in the epidermis is mediated by prostaglandins, presumably PGE_2_. The cell density in the basal layer is approximately 2×10^6^ cells cm^−2^, which is markedly higher than that in MDCK cells (Fig. 2D, 2E). It is not clear whether this may be related to the lower frequency (~300 cm^−2^ h^−1^) and smaller radius of RSPA in the basal layer cells compared to MDCK cells (Fig. 2E).

**Figure 2:**
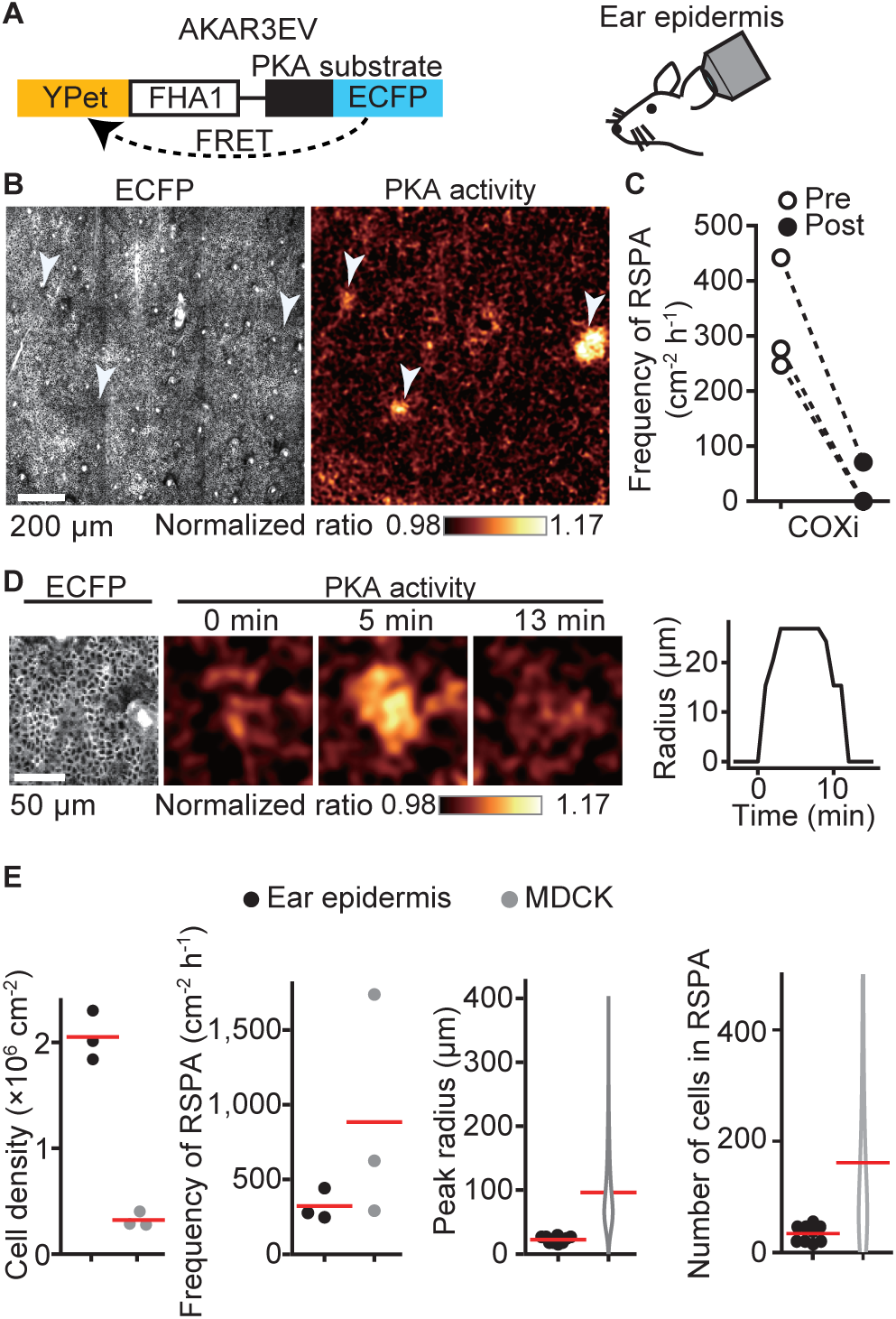
RSPA in the basal layer of mouse auricular epidermis. **(A)** A scheme of AKAR3EV, a PKA sensor. **(B)** Transgenic mice expressing AKAR3EV were observed under a two-photon excitation microscope. Shown are an ECFP image and a YPet/ECFP ratio image representing the cell density and PKA activity in pseudocolor, respectively. (**C**) Three mice were administrated a COXi, 30 mg/kg flurbiprofen intraperitoneally. The frequency of RSPA in pre-treatment was quantitated more than 40 min before the injection. Similarly, the frequency of a post-COX inhibitor treatment was analyzed 15–58, 15–97, and 15–63 min. Each dotted line represents an individual mouse experiment. **(D)** Magnified views of a RSPA *in vivo.* Shown are an ECFP image and a YPet/ECFP ratio image representing the cell density and PKA activity in pseudocolor, respectively. The radius was determined as in Fig. 1C with the detection limit of 6.4 µm. **(E)** The properties of RSPA are compared between *in vivo* and *in vitro*. Data from three independent experiments are shown for each condition. The data of MDCK is from Fig. 1E, seeded at the 1.2 × 10^5^ cells cm^−2^. Cell density and frequency of RSPA data are values per experiment. Peak radius data are pooled from three independent experiments for each condition. Red lines indicate average values.

### RSPA is triggered by calcium transients

What causes RSPA? In agreement with the principal role of calcium in cPLA2 activation, dual imaging of calcium and PKA showed that an intracellular calcium transient precedes RSPA (Fig. 3A). As anticipated, the frequency of RSPA was suppressed by the calcium chelator BAPTA-AM (Fig. 3B). Note that not all of the calcium transients induced RSPA (Fig.3C, arrowheads). Approximately only one-tenth of calcium transients evoke RSPA (compare Fig. 1E and 3D). Moreover, the frequency of calcium transients was also cell density-dependent (Fig. 3D). To further pursue the relationship between calcium transient and RSPA, we employed the Gq-DREADD system (Armbruster et al., 2007). The Gq-DREADD-expressing producer cells were plated with the reporter cells expressing Booster-PKA at a 1:1,200 ratio (Fig 3E). Upon activation of the Gq protein-coupled receptor by the DREADD ligand CNO, we observed RSPA with almost the same size and time course as observed under the non-stimulated condition (Fig. 3F). Among cells with the calcium transient, 76% exhibited RSPA, significantly higher than in the unstimulated state. Because the average calcium signal intensity was higher in RSPA (+) cells than in RSPA (-) cells (Fig. 3G), the peak value of calcium transients appears to be important for the following induction of RSPA.

**Figure 3:**
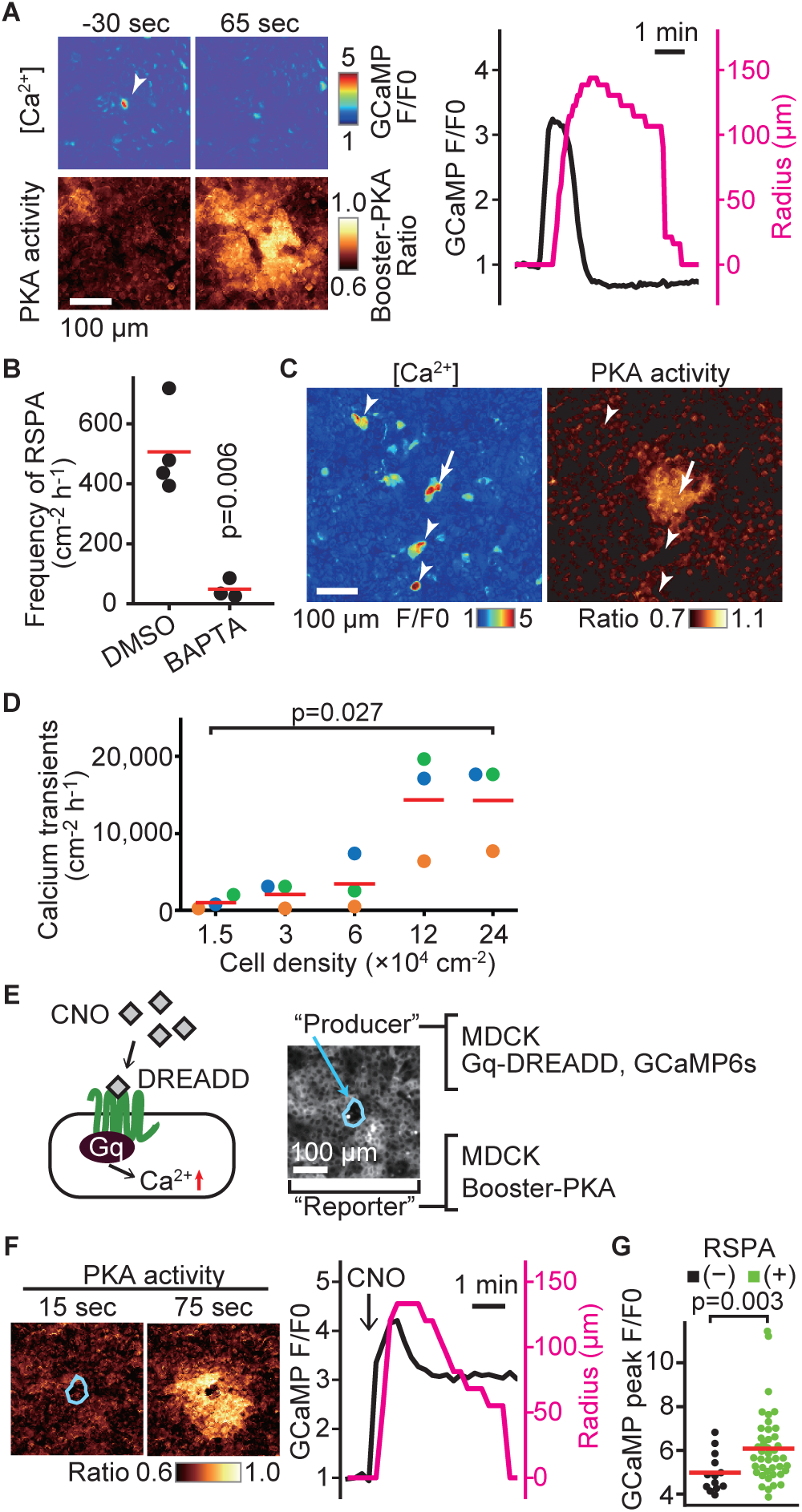
RSPA induction by calcium transients. **(A)** MDCK cells expressing GCaMP6s and Booster-PKA were observed for calcium transients and RSPA. Calcium transients were represented by the fluorescence of GCaMP normalized to the basal level (F/F0). RSPA is analyzed as in Fig. 1C. **(B)** The frequency of RSPA was analyzed 20–80 min after 30 μM BAPTA-AM treatment. Data from three or four independent experiments. The DMSO control data set is from Fig. 1F. **(C)** Maximum projection images of the ratio over 10 min. An arrow or arrowheads represent calcium transients with or without RSPA, respectively. Shown is a part of Video 3. **(D)** MDCK cells expressing GCaMP6s were seeded at the indicated density. Calcium transients showing F/F0 to be greater than three were counted in an indicated cell density. **(E)** Schematic representation of RSPA induction using Gq-DREADD. MDCK cells expressing Gq-DREADD served as producer cells, while MDCK cells expressing Booster-PKA were employed as reporter cells. **(F, G)** MDCK cells expressing Gq-DREADD with GCaMP6s or Booster-PKA were mixed and plated, treated with 1 μM CNO, and imaged. Blue circled cells are producer cells, expressing Gq-DREADD. The FRET ratio, the value of mKate2/mKOκ, in each pixel is shown in pseudocolor as indicated. The time 0 was set as just before CNO addition. In panel I, cells showing an F/F0 value greater than 4 were analyzed for their RSPA as Fig. 1C. Red lines indicate their average value.

### RSPA is a switch-like response to cytoplasmic calcium concentration

To further explore the relationship between the peak value of the calcium transient and RSPA, we employed OptoSTIM1, an optogenetic tool to activate the calcium influx (Kyung et al., 2015)(Fig. 4A). As anticipated, a blue light flash caused a calcium transient, followed by RSPA (Fig. 4B). In a preliminary experiment, a 30 min interval was sufficient to restore the calcium response; therefore, we repeated the blue light flash every 30 min to 1 hour with increasing light intensity. As the light intensity was increased, the amplitude of calcium transients represented by the R-GECO signal also increased linearly (Fig. 4C). In stark contrast, RSPA occurred in an all-or-nothing manner. We repeated this experiment for 13 cells to find any correlations between the calcium concentration and the size of RSPA (Fig. 4D). The R-GECO signal intensity ratio (F/F0) that evoked RSPA ranged from 1.5 to 2.1 among the different cells (Fig. 4D, left), but the size of RSAP did not show a clear correlation with the R-GECO signal intensity ratio (Fig. 4D, right). This observation indicates that there is a threshold of the cytoplasmic calcium concentration for the triggered PGE_2_ secretion.

**Figure 4:**
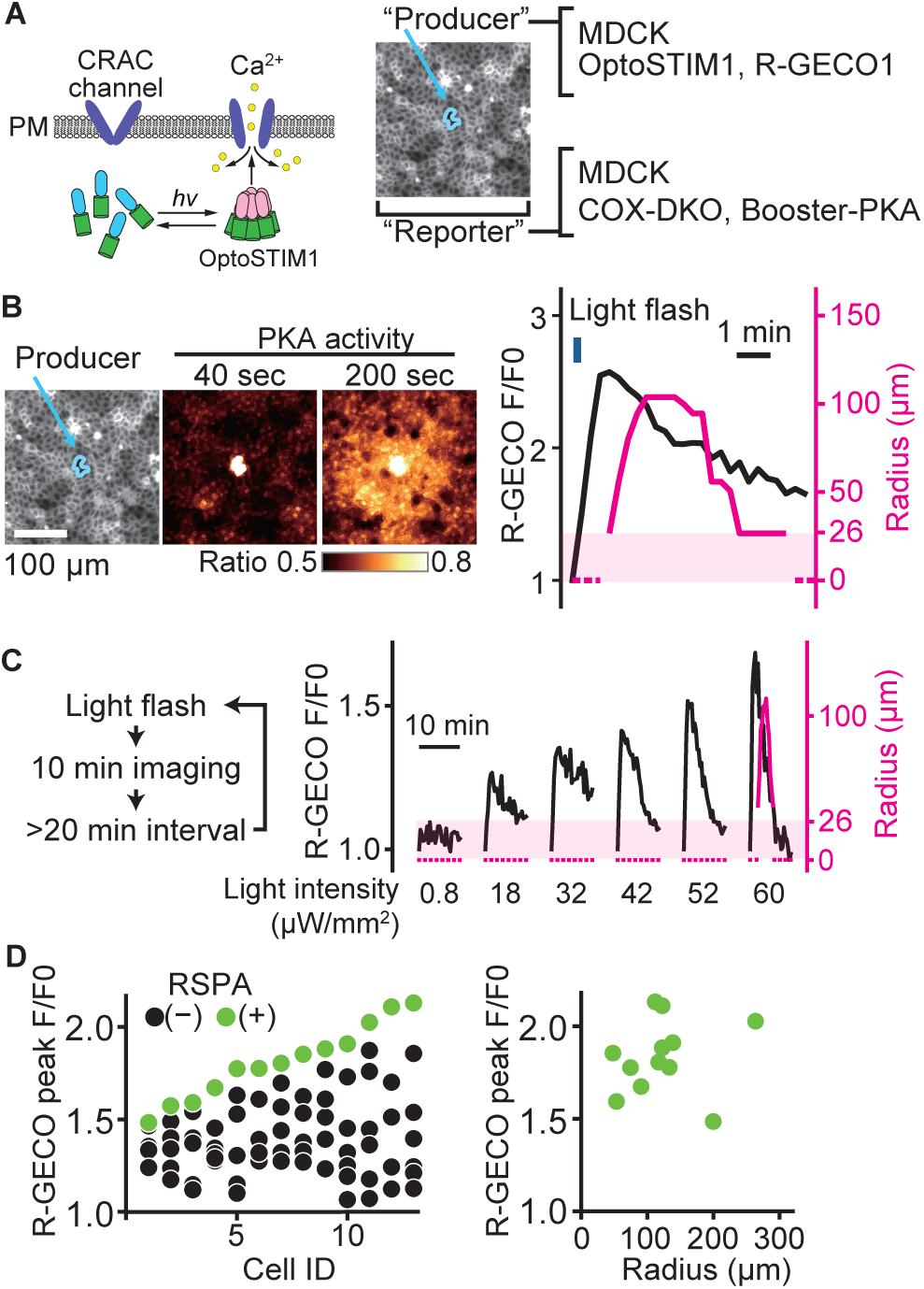
A switch-like response of RSPA to calcium transients. **(A)** Schematic representation of RSPA induction using OptoSTIM1. MDCK cells expressing OptoSTIM1 and R-GECO1 were employed as the producer cells. The Booster-PKA-expressing MDCK cells, deficient in COX-1 and COX-2 (COX-DKO), were employed as the reporter cells. **(B)** The producer cells were stimulated by a flashlight during imaging. Blue circled cells in the mKOκ image are the producer cells. The FRET ratio, the value of mKate2/mKOκ, in each pixel is shown in pseudocolor as indicated. The time 0 was set as just before blue light irradiation. The detection limit for the RSPA radius was 26 µm, as shown in the shaded area. **(C)** Flashlight illumination was repeated with increasing LED power. **(D)** Vertically aligned dots (left) are the results from an individual producer cell. The right panel shows the relationship between R-GECO fluorescence intensity and the radius of RSPA.

### High cell density increases the sensitivity to PGE_2_

We next explored the mechanism that determines the size of RSPA. First, taking advantage of the reproducibility of DREADD system, we examined the involvement of EP2/EP4, cPLA2, and COX1/2 in the calcium-induced RSPA by CRISPR/Cas9-mediated gene knockout (Fig. 5A). As anticipated, we did not observe any RSPA by using the reporter MDCK cells deficient from EP2 and EP4. Knockout of COX2 and cPLA2, but not COX1, in the producer cells almost completely abolished CNO-induced RSPA. These results support the idea that RSPA is mediated by PGE_2_ via the Ca^2+^-cPLA2-COX2-EP2/4-cAMP-PKA pathway (Fig. 5B).

**Figure 5:**
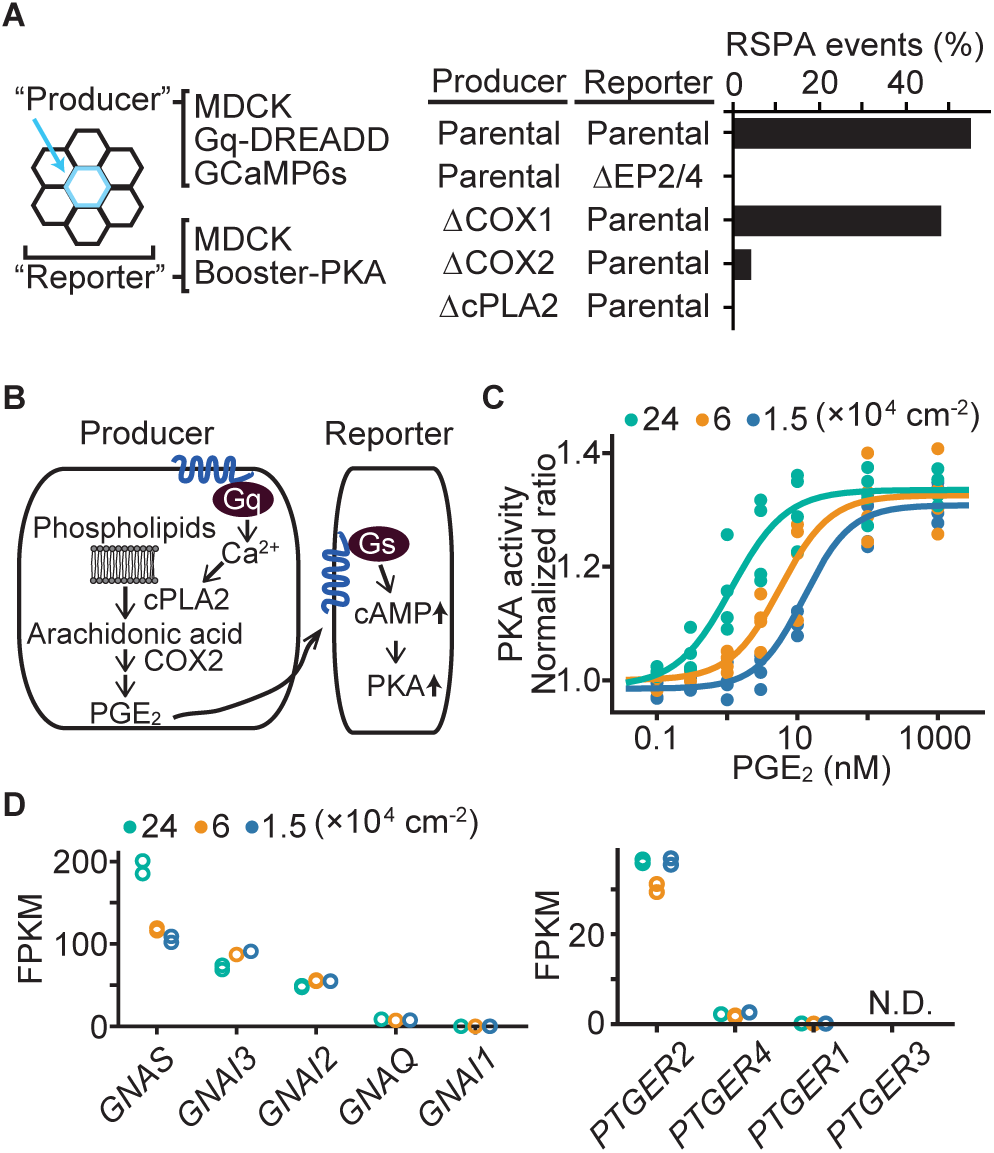
Effect of cell density on PGE_2_ sensitivity and the transcriptome. (**A**) MDCK cells expressing Gq-DREADD and GCaMP6s were employed as the parental producer cells. Meanwhile, MDCK cells expressing Booster-PKA were employed as the parental reporter cells. The genes knocked out by CRIPSR/Cas9 system are depicted in the figure. Analysis was performed as in Fig. 3G. Each producer cell exhibiting F/F0 values greater than 3 was analyzed for the occurrence of RSPA. Data from two independent experiments was summed up. (**B**) Inter- and intra-cellular pathway of RSPA. (**C**) COX-DKO MDCK cells expressing Booster-PKA were plated at the indicated cell density, treated with increasing concentrations of PGE_2_, and analyzed for PKA activity. The mKate2/mKOκ ratio representing PKA activity was calculated and plotted against PGE_2_ concentration. The average intensity of the whole view field of mKate2 or mKOκ, at 20 to 30 min after the addition of PGE_2_, was applied to calculate the mKate2/mKOκ ratio. Three or four independent experiments were performed. (**D**) COX-DKO MDCK cells were seeded at the indicated cell densities and subjected to RNA-Seq analysis. FPKM values of other genes are in Figure S2. N.D. represents Not Detected.

Next, because the size of RSPA depends on the cell density (Fig. 1E), we reasoned that the sensitivity of MDCK cells to PGE_2_ may also be regulated by cell density. To avoid the effect of PGE_2_ produced by the cells, the COX1/2-deficient MDCK cells were challenged by the bath application of PGE_2_. We found an approximately 10-fold difference in the EC50 between the high and low cell densities (Fig. 5C), suggesting that increased sensitivity to PGE_2_ underlies the increased RSPA size under the confluent condition. Transcriptome analysis showed a 2-fold increase in guanine nucleotide-binding protein G(s) subunit alpha (GNAS) (Fig. 5D), but it is not clear whether this difference is sufficient to explain the difference in RSPA frequency. We did not observe any cell density dependence in the transcription of EP2, EP4 (Fig. 5D), phosphodiesterases, or adenylyl cyclases (Fig. S2). Thus, the cell density appears to increase the sensitivity to PGE_2_ mostly in a transcription-independent manner.

### ERK activity is required for RSPA

Previously, we reported that ERK activation is propagated among confluent MDCK cells in a wave-like fashion (Aoki et al., 2013). To examine whether ERK activity also regulates RSPA, we simultaneously observed ERK and PKA activities by using EKAREV-NLS and Boobser-PKA, respectively (Fig. 6A). It appeared that the center of RSPA was localized primarily in areas of high ERK activity. Further quantitative analysis has shown that the ERK activity of the cells locating in the center of RSPA was significantly higher than that of the randomly chosen cells (Fig. 6B, left). However, the size of RSPA did not correlate with the ERK activity (Fig. 6B, right). We next examined the timing of RSPA and the passage of ERK activation waves by aligning the events at the highest PKA activity. It appears that RSPA was evoked when the cells exhibited the highest ERK activity (Fig. 6C, 6D, S4). Cross-correlation analysis of PKA activity and ERK activity revealed that ERK activation preceded PKA activation by approximately 3 min (Fig. 6E). The ERK activation wave is known to be mediated by EGFR and EGFR ligands (Lin et al., 2022). Accordingly, the addition of EGF faintly increased the frequency of RSPA in our experiments, while the MEK and EGFR inhibitors almost completely abrogated RSPA (Fig. 6F), representing that ERK activation or basal ERK activity is essential for RSPA. Collectively, these results obtained with MDCK cells showed that RSPA is triggered by calcium transient in cells with high ERK activity.

**Figure 6:**
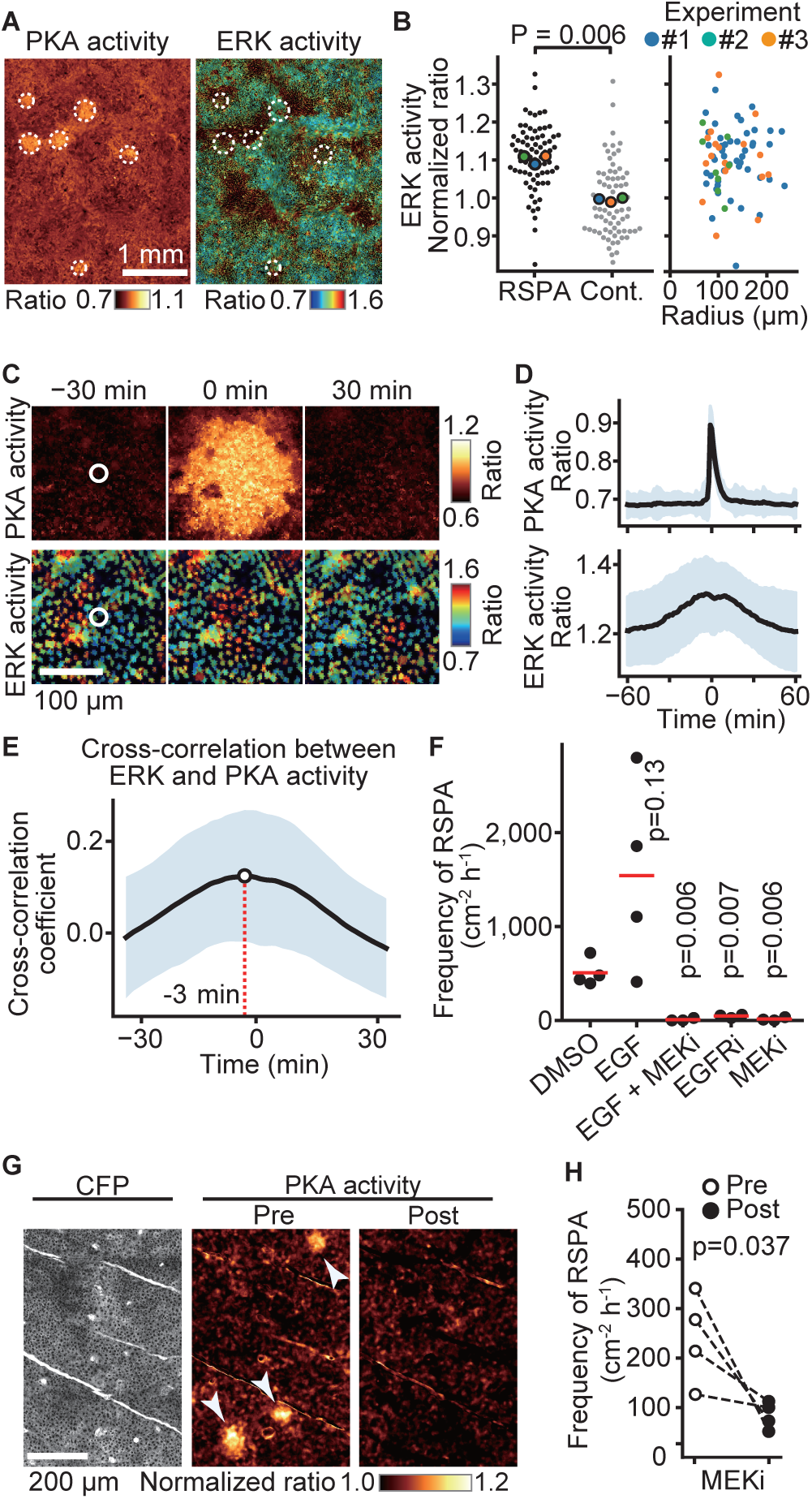
Requirement of ERK activation for RSPA. **(A)** MDCK cells expressing EKAREV and Booster-PKA were observed for ERK and PKA activities every 5 min and 1 min, respectively. The mKate2/mKOκ ratio image represents PKA activity in pseudocolor. The YPet/ECFP ratio image represents ERK activity in IMD mode. RSPA is indicated by white circles. **(B)** The ERK activities within 10 µm from the center of RSPA and within 10 µm from randomly set positions with a random number table generated by Python are plotted in the left panel. Each colored dot represents an average value of an independent experiment. The right scattered plot shows the relationship between ERK activity and the size of RSPA. **(C)** The correlation between ERK activation and RSPA is shown. This is a part of Video 4. **(D)** Cross-correlation analysis of PKA and ERK activities. The average and S. D. values from 67 samples are shown in the black lines and blue shades, respectively. **(E)** Temporal cross-correlations between RSPA and ERK activation rate. The black line indicates the average temporal cross-correlation coefficients with S.D. **(F)** MDCK cells expressing Booster-PKA were imaged in the presence of the following reagents: 0.1% v/v DMSO, 50 ng mL^−1^ EGF, 1 μM PD0325901 (MEKi), and 1 μM AG1478 (EGFRi). The frequency of RSPA was analyzed 20–80 min after the treatment. Each dot represents an individual experiment. Red lines indicate their average value. The control data set is from Fig. 1F. p values were calculated between the labeled sample and the DMSO-treated sample. (**G, H**) Similar to Fig. 2. Transgenic mice expressing AKAR3EV were observed under a two-photon excitation microscope and administrated a MEKi, 5 mg/kg PD0325901 intravenously. Shown are an ECFP image and a YPet/ECFP ratio image representing the cell density and PKA activity in pseudocolor, respectively. The images of PKA activity were projected over 30 min. The frequency of RSPA in pre-treatment was quantitated more than 120 min before the injection. Similarly, the frequency of RSPA in a post-treatment was analyzed 15–90, 15–125, 15-85, and 15–115 min. Each dotted line represents an individual mouse experiment.

The results in MDCK cells motivated us to validate our model *in vivo*. Thus, we tested RSPA in the basal layer of the mouse auricular epidermis could be canceled by the administration of MEKi (Fig. 6G, 6H). As anticipated, RSPA in the basal layer was significantly attenuated 30 minutes after the administration, representing that ERK activity is required for RSPA *in vivo*.

## Discussion

### PGE_2_ discharge causes radial spread of PKA activation in neighboring cells

To the best of our knowledge, only one study has visualized PGE_2_ secretion from a single cell (Zonta et al., 2003). In this study, HEK cells expressing the Gq-coupled PGE_2_ receptor EP1 and a calcium indicator were used to monitor PGE_2_ release from agonist-stimulated astrocytes. However, the spontaneous release of PGE_2_ has never been visualized in either tissue culture cells or live animals. Here we have shown that PGE_2_ is discharged after calcium transients in MDCK cells at high cell densities (Fig. 1). This PGE_2_ discharge leads to the radial spread of PKA activation, which we named RSPA, in the neighboring EP2-expressing cells. By using transgenic mice expressing the PKA biosensor, RSPA was also observed in the mouse auricular epidermis (Fig.2). RSPA in the epidermis is almost completely shut off by COXi, strongly suggesting that prostaglandin(s), most likely PGE_2_, mediates RSPA in the skin. Notably, we failed to observe RSPA in melanoma tissues in which calcium transients were frequently observed (Konishi et al., 2021). We reasoned that repetitive PGE_2_ secretion from tumor cells maintains a high PGE_2_ concentration in the tumor microenvironment, which prevented us from observing pulsatile PKA activation. In fact, the PGE_2_ concentration in melanoma tissue is known to reach as high as 10 µM (Konishi et al., 2021). Thus, RSPA in the skin may function as an alert signal in an early phase of cellular stress.

### PGE_2_ is discharged in a switch-like manner in response to Ca^2+^ transients

Soon after the identification of a Ca^2+^-dependent translocation domain within cPLA2 (Clark et al., 1991), Ca^2+^-dependent cPLA2 arachidonic acid release from cells has been reported (Gijón et al., 2000; Hirabayashi et al., 1999); therefore, it is not surprising to find that the PGE_2_ discharge in our present experiments was due to Ca^2+^-dependent cPLA2 activation (Fig. 3). However, visualization of PGE_2_ secretion at the single-cell resolution revealed a switch-like response of PGE_2_ discharge to the increasing Ca^2+^ concentration (Fig. 4). Recruitment of cPLA2 to the ER and perinuclear membrane requires a higher Ca^2+^ concentration than that to Golgi (Evans et al., 2001). If so, cPLA2 may be sequestered at Golgi at low intracellular Ca^2+^ concentration, and, only when the intracellular Ca^2+^ concentration exceeds the threshold, cPLA2 may reach the ER to liberate arachidonic acids.

### ERK also regulates the probability of RSPA

In the cell density-dependent RSPA of MDCK cells, ERK activity regulates the probability, but not the size, of RSPA. Of note, the MEK inhibitor did not significantly decrease the frequency of calcium transients (Fig. S3), suggesting that ERK had a direct effect on the production of PGE_2_. Because ERK positively regulates cPLA2 by phosphorylating Ser^505^ (Cook & McCormick, 1993; Qiu et al., 1993), it is reasonable that the RSPA is regulated by ERK activity.

The ERK activation wave is operated by a positive feedforward mechanism in which ERK promotes EGFR ligand shedding and the following EGFR activation increases ERK activity in the neighboring cells (Aoki et al., 2017; Hino et al., 2020). Here we found that the ERK activation functions as the “AND gate” for PGE_2_ production together with the calcium transients (Fig. 7). Notably, the propagation of PKA activation, ~100 µm/min (Fig. 1B), is markedly faster than that of ERK activation, 2–4 µm/min (Hiratsuka et al., 2015). Because PKA antagonizes Ras-dependent ERK activation (Burgering et al., 1993; Cook & McCormick, 1993; Wu et al., 1993), the EGFR ligand-ERK and PGE_2_-PKA pathways fit the Turing diffusion reaction model consisting of slow positive and fast negative signaling cascades.

**Figure 7:**
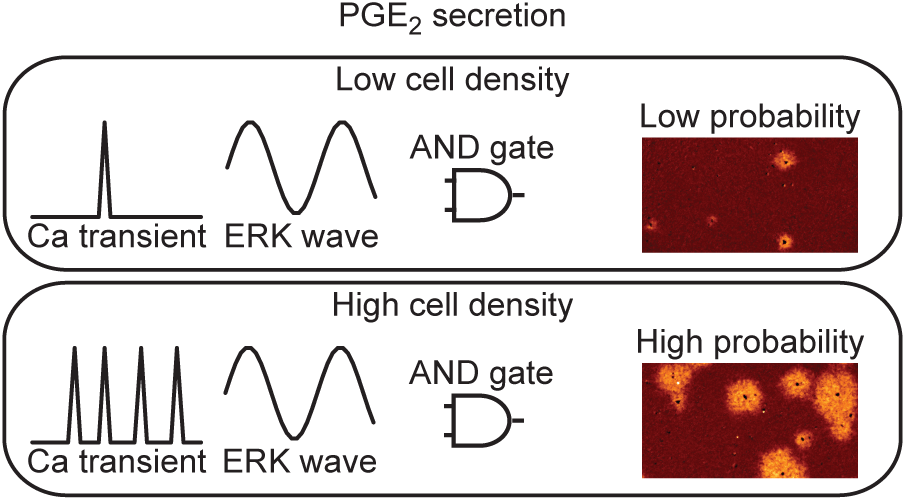
Models for PGE2 secretion. The frequency of calcium transients is cell density-dependent manner. While the ERK activation wave is there in both conditions. Because both calcium transient and ERK activation are required for RSPA, the probability for PGE_2_ secretion is regulated as “AND gate”.

### RSPA may not directly affect cell competition

Recently, PGE_2_ was shown to regulate cell competition among MDCK cells. Interestingly, extrusion of Ras-transformed MDCK cells has been shown to be suppressed by PGE_2_ (Sato et al., 2020), whereas extrusion of MDCK cells expressing constitutively active YAP was dependent on PGE_2_ (Ishihara et al., 2020). Therefore, the effect of PGE_2_ in cell competition could be markedly different according to the signaling cascades that cause the oncogenic changes of MDCK cells.

Importantly, PGE_2_ promotes the extrusion of MDCK cells expressing the constitutively active YAP by internalization of E-cadherin (Ishihara et al., 2020), which is a relatively slow process. Since RSPA causes PKA activation for only several minutes in each cell, multiple RSPA events may be needed to reach the concentration required for the induction of extrusion.

## Conclusions

We have shown that the PGE_2_ discharge from a single cell is a stochastic and switch-like event in the confluent MDCK cells and mouse epidermis. The secreted PGE_2_ can transiently activate PKA in cells within a few hundred micrometers from the producer cell. The question of why cells adopt this pulsatile rather than continuous secretion of PGE_2_ awaits the future.

## Experimental Section

### Reagents

ONO-AE3-208, PF-04418948, clozapine N-oxide, and prostaglandin E_2_ were purchased from Cayman Chemical. BAPTA-AM was obtained from Enzo Life Sciences. PD0325901, mitomycin C, and indomethacin were purchased from FUJIFILM Wako Pure Chemical Corp. AG1478 was purchased from BioVision Inc. Apyrase and EGF were obtained from Sigma-Aldrich. H-89 was purchased from Seikagaku Corp. The DREADD ligand, clozapine N-oxide, was purchased from Cayman Chemical. Flurbiprofen axetil was purchased from KAKEN Pharmaceutical.

### Cell culture

MDCK cells were purchased from the RIKEN BioResource Center (no. RCB0995). Lenti-X 293T cells were obtained from Invitrogen. MDCK and Lenti-X 293T cells were maintained in Dulbecco’s modified Eagle medium (DMEM; FUJIFILM Wako Pure Chemical Corp.) containing 10% fetal bovine serum (Sigma-Aldrich) and 1% v/v penicillin−streptomycin (Nacalai Tesque).

### Plasmids and primers

Plasmids and primers are described in Table S1A and S1B.

### Cell lines

For the generation of MDCK cells stably expressing Booster-PKA or the other ectopic proteins, a lentiviral or piggyBac transposon system was employed. To prepare the lentivirus, a lentiCRISPRv2-derived expression plasmid, psPAX2 (Plasmid: no. 12260; Addgene), and pCMV-VSV-G-RSV-Rev (RIKEN BioResource Center) were co-transfected into Lenti-X 293T cells using polyethyleneimine (Polyscience). Virus-containing media were collected at 48 or 72 h after transfection, filtered, and applied to target cells with 10 μg/mL polybrene (Nacalai Tesque). To introduce ectopic genes using a PiggyBac system, pPB plasmids and pCMV-mPBase(neo-) encoding piggyBac transposase were co-transfected into MDCK cells by electroporation with an Amaxa nucleofector (Lonza). Cells were selected with the medium containing the following antibiotics: 10 μg ml^−1^ blasticidin S (FUJIFILM Wako Pure Chemical Corp.), 100 μg ml^−1^ zeocin (InvivoGen), 2.0 μg ml^−1^ puromycin (InvivoGen), or 200 μg ml^−1^ hygromycin (FUJIFILM Wako Pure Chemical Corp.).

MDCK cells expressing EKAREV-NLS were previously described (Kawabata & Matsuda, 2016). The established cell lines are described in Table S1C.

### CRISPR/Cas9-mediated KO cell lines

For CRISPR/Cas9-mediated single or multiple knockouts of genes, sgRNAs targeting the exons were designed using CRISPRdirect (Naito et al., 2015). Oligo DNAs for the sgRNA were cloned into the lentiCRISPRv2 (plasmid no. 52961; Addgene) vector or pX459 (plasmid no. 62988; Addgene) vector. The expression plasmids for sgRNA and Cas9 were introduced into MDCK cells by lentiviral infection or electroporation. For electroporation, pX459-derived plasmids were transfected into MDCK cells using an Amaxa Nucleofector II. Cells were selected with the medium containing the antibiotics depending on the drug-resistance genes. After the selection, genomic DNAs were isolated with SimplePrep reagent (TaKaRa Bio). PCR was performed using KOD FX neo (Toyobo) for amplification with the designed primers, followed by DNA sequencing.

### Wide-field fluorescence microscopy

Cells were imaged with an ECLIPSE Ti2 inverted microscope (Nikon) or an IX83 inverted microscope (Olympus). The ECLIPSE Ti2 inverted microscope was equipped with a Plan Fluor 10X or 4X objective, an ORCA Fusion Digital CMOS camera (HAMAMATSU PHOTONICS K.K.), an X-Cite TURBO LED light source (Excelitas Technologies), a Perfect Focus System (Nikon), a TI2-S-SE-E motorized stage (Nikon), and a stage top incubator (Tokai Hit). The IX83 inverted microscope was equipped with a UPlanAPO 10x/0.40 NA objective lens (Olympus), a Prime sCMOS camera (Photometrics), a CoolLED precisExcite LED illumination system (Molecular Devices), an IX2-ZDC laser-based autofocusing system (Olympus), and an

### MD-XY30100T-Meta automatically programmable XY stage

The following filters were used for the multiplexed imaging: for CFP and YFP imaging, a 434/32 excitation filter (Nikon), a dichroic mirror 455 (Nikon), and 480/40 and 535-30 emission filters (Nikon) for CFP and YFP, respectively; for GCaMP6s imaging, a 480/40 (Nikon) excitation filter, a dichroic mirror 455 (Nikon), and a 535/50 emission filter (Nikon); for mKOκ and mKate2 imaging, a 555BP10 excitation filter (Omega Optical), an FF562Di03 dichroic mirror (Semrock), and XF3024 (590DF35) (Omega Optical) and BLP01-633R-25 (Semrock) emission filters for mKOκ and mKate2, respectively; for iRFP670 imaging, an FF01-640/14 excitation filter (Semrock), a dichroic mirror 660 (Nikon), and a 700/75 emission filter (Nikon); for R-GECO1, a 555BP10 excitation filter (Omega Optical), an FF562Di03 dichroic mirror (Semrock), and an XF3024 emission filter (590DF35) (Omega Optical).

### *In vivo* two-photon imaging of the mouse epidermis

The establishment of transgenic mice expressing AKAR3EV (PKAchu mice) was described previously (Kamioka et al., 2012). Briefly, 8- to 13-week-old female mice were used for the in vivo imaging. The ear hair was removed with a razor one day before the experiments. Mice were anesthetized with 1.5% isoflurane (FUJIFILM Wako Pure Chemical Corp.) inhalation and placed in a side-lying position on an electric heater maintained at 37°C. The ear skin was placed on the cover glass. Two-photon excitation microscopy was performed with an FV1200MPE-IX83 inverted microscope (Olympus) equipped with a 30x/1.05 silicon oil-immersion objective lens (XLPLN 25XWMP; Olympus), an InSight DeepSee Ultrafast laser (Spectra Physics), an IR-cut filter (BA685RIF-3), two dichroic mirrors DM505 (Olympus), and two emission filters (BA460-500 for CFP and BA520-560 for YFP) (Olympus). The excitation wavelength was 840 nm.

### Spontaneous RSPA

MDCK cells expressing Booster-PKA or hyBRET-Epac were seeded on collagen-coated glass-bottom 96-well plates (Matsunami Glass Ind.) at a density of 1.2 to 2.4 × 10^5^ cells/cm^2^. Before imaging, the culture media were replaced with phenol red-free M199 (ThermoFisher Scientific) supplemented with 10% fetal bovine serum. Cells were imaged by wide-field fluorescence microscopy, as described above.

### Analysis of calcium concentrations

Intracellular Ca^2+^ concentrations in MDCK cells were visualized with a genetically encoded calcium indicator, GCaMP6s or R-GECO1. For GCaMP6s analysis, calcium signals were expressed as F/F0, where F is the fluorescence at each time point, and F0 represents baseline fluorescence. To analyze the peak F/F0 value of GCaMP6s, F0 was calculated as the minimum projection of fluorescence intensity over the 5 min before each frame. Each cell showing calcium transient was visually checked to exclude the F/F0 elevation caused by flowing debris and misregistration of cells. If two or more adjacent cells showed calcium transients simultaneously, it was counted as a calcium transient. If two or more calcium transients were detected at intervals of more than one minute, they were counted separately. In the Gq-DREADD experiment, F0 was calculated as the mean intensity before the stimulation.

For R-GECO1 analysis, F/F0 calcium signals were calculated by assigning the reference F0 using the fluorescence intensity before each blue light flash.

### Gq-DREADD-induced calcium transients and RSPA

MDCK cells expressing both Gq-DREADD-P2A-mCherry-NLS (Evans et al., 2001) and GCaMP6s (Evans et al., 2001) were utilized as PGE_2_ producer cells. MDCK cells expressing Booster-PKA served as PGE_2_ reporter cells. The producer and reporter cells were mixed at a ratio of 1:400 to 1:200 and plated on collagen-coated glass-bottom 96-well plates (Matsunami Glass Ind. or AGC Inc.) at a density of 1.2 to 2.4 × 10^5^ cells/cm^2^. Before imaging, the culture media were replaced with phenol red-free M199 (ThermoFisher Scientific) supplemented with 10% fetal bovine serum. Gq-DREADD was activated by the addition of 1 µM of clozapine N-oxide (CNO). Cells were imaged by wide-field fluorescence microscopy, as described above.

### Light-induced calcium transients and RSPA

MDCK cells expressing both R-GECO1-P2A-iRFP670 and OptoSTIM1 (CRY2clust) (Lee et al., 2014) were used as PGE_2_ producer cells. COX1 and COX2-deficient MDCK cells expressing Booster-PKA were used as PGE_2_ reporter cells. The producer and reporter cells were mixed at a ratio of 1:400 to 1:1,200 and plated on collagen-coated glass-bottom 96-well plates (Matsunami Glass Ind.) or 24-well plates (AGC Inc). After 16 to 32 h of incubation, the culture media were replaced with phenol red-free M199 (ThermoFisher Scientific) supplemented with 10% fetal bovine serum or M199 supplemented with 0.1% w/v bovine serum albumin (Sigma-Aldrich). Cells were imaged by wide-field fluorescence microscopy, as described above. During the observation, OptoSTIM1 was activated with 475 nm LED for 200 msec to trigger calcium influx into the cell. To control the calcium influx from small to large, the excitation light was modulated from 0.8 to 67 μW mm^−2^. To prevent cell division, MDCK cells with 3 µg/ml of mitomycin C for 1 h one day before passage.

### Titration of PGE_2_ sensitivity

COX-1 and COX-2 depleted (COX-DKO) MDCK cells expressing Booster-PKA were seeded on a 96-well glass-base plate at the indicated densities. Before imaging, the culture media were replaced with phenol red-free M199 (ThermoFisher Scientific) supplemented with 10% fetal bovine serum for MDCK. The 96-well plate was imaged by an inverted microscope as described earlier. mKate2 and mKOκ images were obtained in one position for every well at around 5 min intervals. Cells were stimulated with PGE_2_ at the indicated concentrations. The mKate2/mKOκ ratio was quantified from the average intensity of the whole field of view at around 20 to 30 min after the addition of PGE_2_.

### Quantification of RSPA

The program code for image analysis is available via GitHub at https://github.com/TetsuyaWatabe-1991/RSPAanalysis.

Ratio images of MDCK cells expressing Booster-PKA were created after background subtraction. A median filter and a Gaussian 2D filter were applied to each image for noise reduction. The ratio image was normalized by a minimum intensity projection along the time axis. The processed images were binarized with a predetermined threshold and processed by morphological opening and closing to refine the RSPA area. Center coordinates and equivalent circle radii were obtained from each RSPA area. If the distance between the center coordinates of RSPA between successive frames was less than 100 μm, they were considered to be the same RSPA. For MDCK cells expressing hyBRET-Epac and mouse ear skin expressing AKAR3EV, the center coordinates of each RSPA were manually determined due to the low signal-to-noise ratio.

To obtain the time course of the RSPA radius, concentric regions were defined at the center of each RSPA. The median FRET ratio in each concentric ROI was calculated. The radius of the outermost concentric region where the median ratio value exceeds a predetermined threshold was defined as the radius of the RSPA.

### Cross-correlation analysis of ERK and PKA activity

Cross-correlation analysis was performed with Python using the scientific library SciPy (http://www.scipy.org). The centers of each spontaneous RSPA in MDCK cells expressing both EKAREV and Booster-PKA were detected automatically as described above. The regions of interest were defined at the center of each RSPA with a radius of 10 µm, and the average ERK and PKA activity from 2 to 5 cells was quantified. The program code for this analysis is available via GitHub as described above.

### RNA-Seq

COX1 and COX2-deficient MDCK cells expressing Booster-PKA were seeded in collagen-coated glass-bottom 96-well plates (AGC Inc.) at a density of 1.5 × 10^4^ or 6.0 × 10^4^ or 2.4 × 10^5^ cells/cm^2^. After 24 h of incubation, the culture media were replaced with phenol red-free M199 (ThermoFisher Scientific) supplemented with 10% fetal bovine serum. Three hours after medium replacement, RNA was extracted from each sample using an RNeasy Mini Kit (Qiagen). Libraries for RNA-Seq were prepared using an NEBNext Ultra II Directional RNA Library Prep Kit for Illumina (New England Biolabs) and sequenced on the NextSeq500 (Illumina) as 75 bp single-end reads. RNA-Seq data were trimmed using Trim Galore version 0.6.6 (Krueger 2015) and Cutadapt version 2.8 (Martin 2011). The quality of reads was checked and filtered using FastQC version 0.11.9 (Andrews 2010). The reads were mapped to a reference genome canFam3.1 (Lindblad-Toh 2005, Hoeppner 2014) using HISAT2 version 2.2.1 (Kim 2019), and the resulting aligned reads were sorted and indexed using SAMtools version 1.7 (Li 2009). Relative abundances of genes were measured in FPKM using StringTie version 2.1.4 (Kovaka 2019, Pertea 2015). Plots were created in Python using the pandas, matplotlib, NumPy, and seaborn libraries.

Sequence data are available in the DNA Data Bank of Japan Sequence Read Archive under accession numbers DRR014156 to DRR014161

### Statistical analysis

All statistical analyses and visualizations were performed in Python using the libraries NumPy, pandas, SciPy, pingouin, matplotlib, and seaborn. No statistical analysis was used to predetermine the sample size. Welch’s t-test was used to evaluate statistically significant diLJerences. The p values less than 0.05 were considered statistically significant.

## Data availability

The data that support the findings of this study are available within the article and its Supplementary Information or from the corresponding author upon reasonable request.

## ACKNOWLEDGMENTS

We are grateful to the members of the Matsuda Laboratory for their helpful input, K. Hirano, T. Uesugi, and Y. Takeshita, who provided technical assistance, and to the Medical Research Support Center of Kyoto University for DNA sequence analysis. We thank Takefumi Kondo and Yukari Sando (NGS core facility of the Graduate Schools of Biostudies, Kyoto University) for supporting the RNA-Seq analysis. This work was supported by the Kyoto University Live Imaging Center. Financial support was provided by JSPS KAKENHI grants (nos. 21K20773 to T.W., 21H02715 and 21H05226 to K.Terai, and 19H00993 and 20H05898 to M.M.), a JST CREST grant (no. JPMJCR1654 to M.M.), a JST Moonshot R&D grant (no. JPMJPS2022 to M.M.), and a grant from Fugaku Foundation (to M.M.).

## AUTHOR INFORMATION

### Corresponding Author

Kenta Terai − Department of Pathology and Biology of Diseases, Graduate School of Medicine, Kyoto University, Kyoto 606-8501, Japan

### Author Contributions

Conceptualization, methodology, validation, formal Analysis: T.W., K.Terai; Investigation: T.W., S.Y., K.Takakura and K.Terai; Data Curation: T.W. and K.Terai; Resources: T.W., K.Terai and M.M.; Writing – Original Draft: T.W.; Writing – Review & Editing: D.T., S.N., K.Terai. and M.M.; Supervision: K.Terai and M.M.; Project Administration: K.Terai and M.M.; Funding Acquisition: T.W., K.Terai and M.M.

### Notes

The authors declare no competing financial interest.

## Supplementary information

**<u>Figure S1: The probability of RSPA in each cell</u>**

**(A)** MDCK cells expressing Booster-PKA were seeded at the indicated density and analyzed. Each color represents an individual experiment. Red lines indicate average values. (**B**) A scheme of hyBRET-Epac, a cAMP sensor. (**C**) MDCK cells expressing hyBRET-Epac were imaged to generate Turquoise/YPet ratio images representing cAMP concentration, [cAMP], in pseudocolor. The image of Turquoise represents the cell density, which is seeded at 1.2 × 10^5^ cells cm^−2^. The time 0 was set as just before cAMP production. The normarized ratio images were binarized with the threshold value 1.06 of Turquoise/YPet ratio. (**D**) MDCK expressing hyBRET-Epac or Booster-PKA were seeded at seeded at 2.4 × 10^5^ cells cm^−2^. The gradients of normalized ratio were measured with a 10 pixel-width line scanning across the center of RSPA. Each color represents an individual RSPA. (**E**) MDCK cells expressing hyBRET-Epac were seeded at 2.4 × 10^5^ cells cm^−2^ and analyzed for the peak radius of RSPA.

**<u>Figure S2: Effect of cell density on the transcriptome</u>**

COX-DKO MDCK cells were seeded at the indicated cell densities and subjected to RNA-Seq analysis. FPKM values of genes related to PGE_2_ homeostasis: isoforms of phosphodiesterase (PDE), isoforms of adenylyl cyclase (ADCY), and phospholipase A2 (PLA2G4A). In agreement with previous reports, Yap target genes, CYR61, CTGF, and AXL, were suppressed at high cell density.

**<u>Figure S3: Effect of MEK inhibitor on calcium transients</u>**

MDCK cells expressing GCaMP6s were seeded at 1.2 × 10^5^ cells cm^−2^. Cells were incubated with 0.1% v/v DMSO or 1 mM PD0325901 (MEKi) for 90 min and imaged every 5 sec for 20 min. Cells in interphase showing the indicated values of F/F0 peak were counted and shown as calcium transients per cell per hour. Data are from a field of view containing around 2.0 × 10^4^ cells from a single experiment.

**<u>Figure S4: Representative ERK and PKA activities in the center of RSPA</u>**

Five representative plots of ERK and PKA activities in Fig. 5D. Each color represents an individual RSPA.

**<u>Video 1: RSPA in MDCK cells.</u>**

The experiments described in Fig. 1B are performed and analyzed.

**<u>Video 2: RSPA in living mice.</u>**

The experiments described in Fig. 2B are performed and analyzed.

**<u>Video 3: Correlation of calcium concentration with RSPA.</u>**

The experiments described in Fig. 3C are performed and analyzed.

**<u>Video 4: Correlation of ERK activity with RSPA.</u>**

The experiments described in Fig. 6C are performed and analyzed.

**<u>Video 5: Requirement of ERK activation for RSPA *in vivo*.</u>**

The experiments described in Fig. 6G are performed and analyzed.

