## Supplementary figures and images for "Calcium transients trigger switch-like discharge of prostaglandin E_2_ in an extracellular signal-regulated kinase-dependent manner"

### Supplemental figures

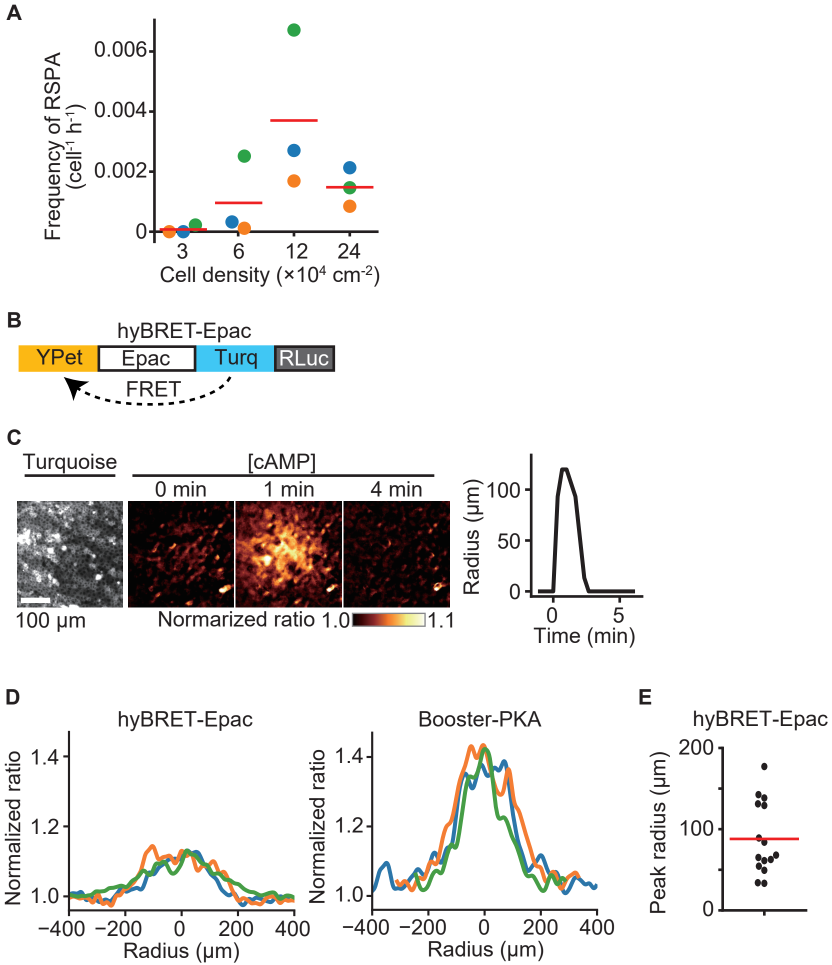


Watabe et al. Fig. S1

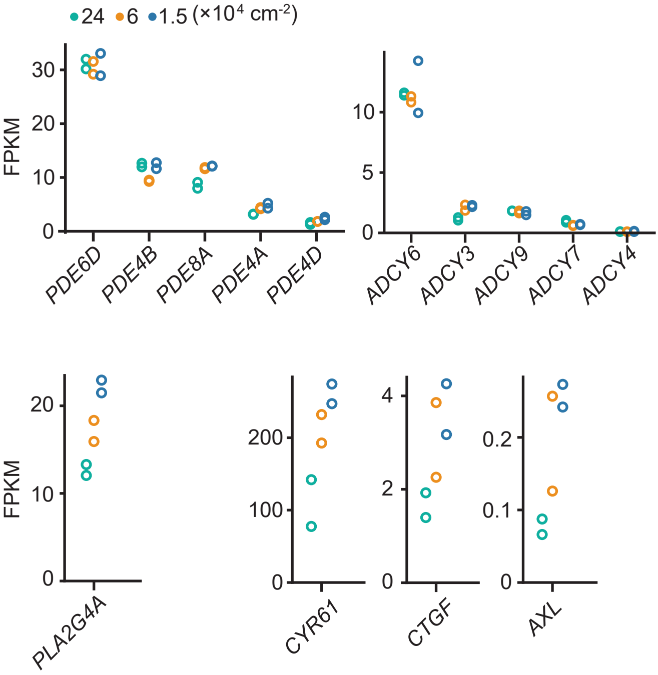


Watabe et al. Fig. S2

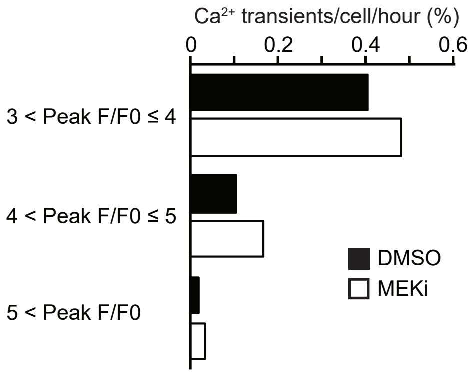


Watabe et al. Fig.S3

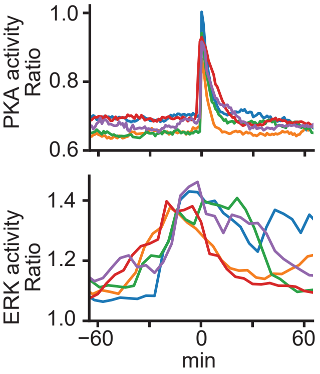


Watabe et al. Fig.S4
